# Medial prefrontal cortex activity reflects ordinal retrieval modes and hippocampal activity reflects temporal context retrieval modes

**DOI:** 10.1101/501122

**Authors:** Puck C. Reeders, Amanda G. Hamm, Timothy A. Allen, Aaron T. Mattfeld

## Abstract

Remembering sequences of events defines episodic memory, but retrieval can be driven by both ordinality and temporal contexts. Whether these modes of retrieval operate at the same time or not remains unclear. Theoretically, medial prefrontal cortex (mPFC) confers ordinality, while the hippocampus (HC) associates events in gradually changing temporal contexts. Here, we looked for evidence of each with BOLD fMRI in a sequence task that taxes both retrieval modes. To test ordinal modes, items were transferred between sequences but retained their position (e.g., AB<u>3</u>). Ordinal modes activated mPFC, but not HC. To test temporal contexts, we examined items that skipped ahead across lag distances (e.g., AB<u>D</u>). HC, but not mPFC, tracked temporal contexts. There was a mPFC and HC by retrieval mode interaction. These current results suggest that the mPFC and HC are concurrently engaged in different retrieval modes in support of remembering *when* an event occurred.

**Significance Statement:** Memory for sequences of events is a defining aspect of everyday episodic memory allowing our brain to separate unique experiences that otherwise have overlapping sensory and spatial content. Sequence memory is impaired in typical aging and in disorders such as Alzheimer’s disease. The results of the current study provide new evidence that two retrieval modes concurrently arise during sequence memory, and they have distinct neural correlates. The medial prefrontal cortex contributes to an ordinal retrieval mode, while at the same time, the hippocampus contributes a gradually-changing temporal context mode of retrieval. These data shed new light on why typical episodic memory requires both the medial prefrontal cortex and the hippocampus, and suggests a functional dissociation between the medial prefrontal cortex and hippocampus across these modes of retrieval.

## Introduction

Memory for sequences of events is a fundamental component of episodic memory (Tulving 1984; 2002; Allen and Fortin, 2013; Howard and Eichenbaum, 2013; Eichenbaum, 2017). While different experiences share overlapping elements, the sequence of events is unique. Remembering the order of events allows us to disambiguate episodes with similar content and make detailed predictions supporting decision making.

At least two complementary memory processes contribute to the retrieval of events in the correct sequence: ordinal (Orlov et al., 2002) and temporal context (Howard and Kahana, 2002) retrieval modes. Whether these disparate retrieval modes operate coincidently or not remains an open question with consequences for understanding basic mechanisms of how we remember the events that unfold throughout our day. According to an ordinal retrieval mode, items are remembered by their position within an event sequence (Dubrow and Davachi, 2013; Allen et al., 2014; Long and Kahana, 2019) providing for sequential memory through well-established semantic or abstracted relationships (1^st^ then 2^nd^, etc.). While for a temporal context retrieval mode, events are remembered through a gradually changing temporal context within which specific items have been associated. According to temporal context, when an element of a sequence is presented or retrieved (e.g., ‘C’ in AB**C**DEF), items that are close in the sequence (e.g. the ‘D’ in the sequence) have a higher retrieval rate compared to items that are further away (e.g. the ‘F’ in the sequence). These temporal contexts result from item associations that are dependent on time varying neural activity (e.g., Eichenbaum, 2014), and contribute to sequence memory through the reactivation of neighboring items during retrieval (Dubrow and Davachi, 2013; Long and Kahana, 2019).

The medial prefrontal cortex (mPFC) and hippocampus (HC) are thought to contribute to sequence memory through ordinal representations and temporal contexts, respectively (Agster et al., 2002; Fortin et al., 2002; Kesner et al., 2002; DeVito and Eichenbaum, 2011; Allen et al., 2016; Jenkins and Ranganath, 2016). In rodents, mPFC disruptions impair sequence memory (DeVito et al., 2011; Jayachandran et al., 2019), mPFC ‘time cells’ are evident (Tiganj et al., 2017), and positions within a sequence can be the main determinant of differential activity in mPFC neurons during spatial sequences (Euston and McNaughton, 2006). In humans, mPFC activation is sensitive to temporal order memory (Preston and Eichenbaum, 2013), and codes for information about temporal positions within image sequences irrespective of the image itself (Hsieh and Ranganath, 2015). HC activations are also generally associated with temporal order memory (Kumaran and Maguire, 2006; Ekstrom and Bookheimer, 2007; Lehn et al., 2009; Ross et al., 2009; Jenkins and Ranganath, 2010; Tubridy and Davachi, 2011; Kalm et al., 2013; Hsieh et al., 2014; Goyal et al., 2018). The HC binds events within temporal contexts (Eichenbaum et al., 2007; Dubrow and Davachi, 2013; Bladon et al., 2019) through a gradually changing neural context (Manns et al., 2007; Mankin et al., 2012). Similarly, medial temporal lobe neuronal and BOLD activations in humans have demonstrated evidence for gradually evolving temporal contexts (Howard et al., 2012; Kalm et al., 2013; Kragel et al., 2015).

Here we tested the contributions of the mPFC and HC during a visual sequence memory task that provides behavioral evidence of both ordinal and temporal context retrieval modes (Allen et al., 2014). In the task, these two retrieval modes are parsed using probe trials that place conflicting demands on ordinal (Allen et al., 2014; 2015; Orlov et al., 2000) and temporal context modes (Jayachandran et al., 2019). We first evaluated ordinal retrieval modes using items that were transferred from one sequence to another while retaining their ordinal position. Evidence for an ordinal-based retrieval mode occurs when these probes are identified as in sequence because they occur in the same ordinal position of their original sequence. mPFC activations (but not HC) was strongest for these ordinal retrievals, and did not reflect general task accuracy. Second, we evaluated a temporal context retrieval mode using items that skipped ahead with shorter lag distances (ABC**F**EF) to larger lag distances away from their position (A**F**CDEF). HC activations (but not mPFC) tracked with lag distance, providing evidence the HC reflects a temporal context-based retrieval mode. In fact, a significant interaction was observed such that mPFC and HC differentially activated for ordinal and temporal context retrievals. Our data shows that sequence memory involves both retrieval modes. In line with these results, we suggest that understanding episodic memory requires more insight into the neurobiology of ordinal processing in mPFC, in addition to the more often studied temporal contexts in HC.

## Materials and Methods

### Participants

Thirty-nine right-handed volunteers were recruited from Florida International University (FIU) and University of Miami to perform a magnetic resonance imaging (MRI) study that included a sequence memory task designed to investigate the ability of humans to learn and remember arbitrary sequences of items, and a temporal reward discounting task. Task order was counterbalanced across all participants. Here we report results from the sequence memory task. All participants provided written consent in compliance with the local Institutional Review Board. Five participants were excluded from the final analysis due to failure to complete the task (n = 1) or because of poor performance (n = 4; sequence memory index score ≤ 0; see Sequence Memory Analysis section below). Participants excluded for poor performance had d-prime scores 2 standard deviations below the mean (M = 2.030, SD = 0.807). The final sample consisted of 34 individuals (19 females; mean age = 21 years, SD = 2).

### Task Apparatus

The sequence task was run on a Dell computer using Matlab (R2015b) with custom scripts that included functions from Psychtoolbox (Psychtoolbox-3 distribution; http://www.psychtoolbox.org). Images were back-projected and viewed by participants with an angled mirror mounted on the head coil. Responses were recorded using a Current Designs MR-compatible 4-button inline response device (https://www.curdes.com).

### Experimental Design and Statistical Analysis

#### Prescan Training

All participants began with a practice session to become acquainted with the structure of the task and the method of responding. During the practice session, participants viewed four low-memory demand sequence sets, each comprising six unique images: (1) individual arrows at 0°, 60°, 120°, 180°, 240°, and 300°, presented in a clockwise fashion (Fig1A, Seq1); (2) a dot moving from the upper left to the lower right corner (Fig. 1A, Seq2); (3) bars of different colors moving from left to right (not shown); and (4) letters A, B, C, D, E, and F positioned in the center of the screen (not shown). Participants were asked to memorize sequences after a single “study” presentation during which the items were passively viewed. Later the practice sequences were tested with 15 unique memory probes per sequence. Testing was self-paced; each sequence was preceded by a screen with the words “Press the button to begin”. To initiate an image in the sequence, participants were required to press and hold a button. If the image was in sequence (InSeq), participants were instructed to hold down the button until the image disappeared on its own at 1 s (the decision threshold), after which they could release the button. If the item was out of sequence (OutSeq), participants were instructed to release the button prior to the decision threshold (<1 s), at which point the image would disappear upon button release. Prescan training was conducted on a Dell desktop computer.

**Figure 1.**
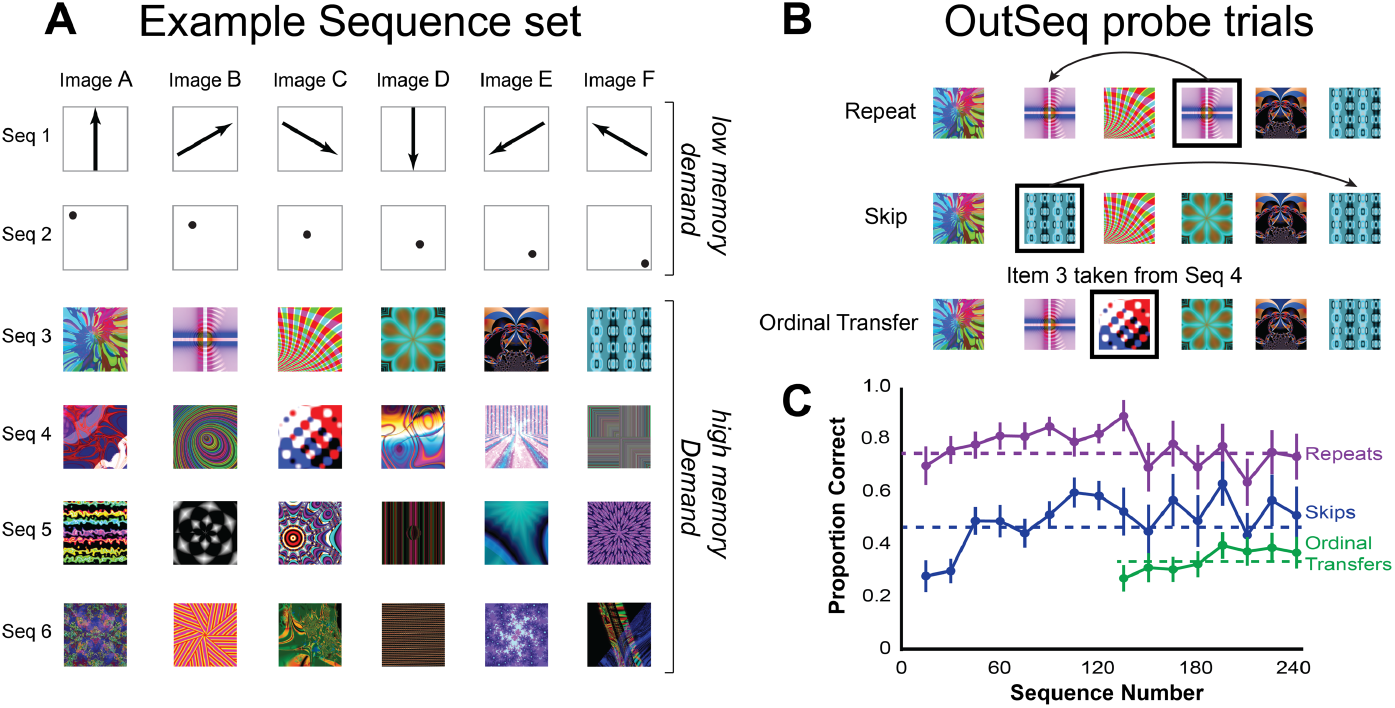
Sequence memory task and overall performance levels. Participants were tested on a sequence memory task that pressures different retrieval modes using different out of sequence probe trial types. **A,** participants were presented sequences, one item at a time, to study before being tested on whether an item was in sequence or out of sequence. The total sequence set included 6 sequences, from which two sequences were *low memory demand* sequences (Seq 1-2), which were easy, predictable sequences that could be solved using a rule-based strategy. The other four sequence were *high memory demand* sequences (Seq 3-6), that required memorization of the items during the study phase. **B,** there were three out of sequence probe trials: Items that were repeated in the sequence (Repeats), items that were presented too early in the sequence (Skips), and items that transferred from one sequence to another, while remaining in their ordinal position (Ordinal Transfers). Repeats and Skips occurred throughout the whole task, whereas Ordinal Transfers occurred during the second half to avoid participants relying solely on ordinal strategies. **C,** participants performed best on identifying Repeats, then Skips and then Ordinal Transfers. For all three probe trial types, learning was rapid (asymptotic within a few trials), and performance was steady throughout the experiment. Thus, performance in the task and the accompanying brain imaging data primarily reflect memory and the associated retrieval modes.

#### Sequence Memory Task

During the Sequence Memory Task, participants were presented with two sequences from prescan training (*low-memory demand;* Fig. 1A, Seq 1-2), and four novel sequences consisting of six unique fractal images (*high-memory demand;* Fig. 1A, Seq 3-6). The exact composition of the novel fractal sequence sets was different for each participant. Sets were selected randomly, without replacement, from a bank of 240 unique fractal images. Similar to the prescan training, participants were asked to view and memorize the sequences after a single presentation in a passive viewing phase. Following the passive viewing phase, sequences were presented in a pseudorandom order, participants made judgments as to whether each item in a sequence was presented InSeq or OutSeq.

Testing was self-paced and followed the same structure as the prescan training. The words “Press the button to begin” preceded the beginning of each sequence. A button depression with the right index finger initiated the presentation of each image in a sequence. If the participants judged the current image to be InSeq, they would hold the button until the decision threshold (1 s, when the image disappeared), after which they could release the button. If the image was OutSeq, they were instructed to release the button before the 1 s decision threshold. Sequence order was determined by the following rules: (1) each sequence was presented first with all images in the correct order. (2) In the first half of testing (first 120 sequences), OutSeq items were either the same image appearing twice (Repeats) or an item appearing too early (Skips; see probe trial description below). In the second half of testing (second 120 sequences), OutSeq items could now include images appearing from a different sequence, but in the correct ordinal position (Ordinal Transfers). Ordinal Transfers were introduced later so participants would not adopt an explicit ordinality strategy at the outset of the study given the dominance of ordinal retrieval modes early in sequence memory (Orlov et al., 2000). The six sequences were presented 40 times each, for a total of 240 sequence presentations. Half of the total sequences contained one OutSeq and five InSeq images; the remaining consisted only of correctly sequenced items. InSeq and OutSeq sequence sets were randomly presented throughout testing. The Sequence Memory Task consisted of 15 min blocks of continuous performance separated by a brief break (<1 min) to provide participants a rest. The number of blocks was dependent on the pace of the participant. Out of all the analyzed participants, most participants completed the task in four blocks (n = 29), while the remainder finished in either three (n=1) or five (n=4).

#### OutSeq Probe Trials

Three distinct types of OutSeq probe trials were used during the test phase: Ordinal Transfers, Repeats (reverse lags) and Skips (forward lags). OutSeq items were counterbalanced across sequence sets, never presented in first position (Pos1), and each OutSeq instance was unique. Due to the nature of the OutSeq trials, there could not be equal distribution of probe trials across all possible positions, and occurred as follows: Pos1, 0%; Pos2, 7.5%; Pos3, 17,5%; Pos4, 50%; Pos5, 17.5%; Pos6, 7.5%.

*Ordinal Transfers*. OutSeq trials where an item from one sequence was ‘transferred’ to a different sequence while retaining its correct ordinal position were considered Ordinal Transfers (Fig. 1B, *Ordinal Transfer*). For example, consider two sequences consisting of ABCDEF and UVWXYZ. An ordinal transfer into the first sequence might have Y at the 5^th^ position ABCD<u>Y</u>F. While Y occupies its original 5^th^ position, it otherwise does not belong to the current sequence (Y does not normally follow D). Distribution of Ordinal Transfers across all possible positions: Pos1, 0%; Pos2, 7.5%; Pos3, 17,5%; Pos4, 50%; Pos5, 17.5%; Pos6, 7.5%.

*Repeats (reverse lags).* An OutSeq item was considered a Repeat if the image previously appeared in the current sequence and was repeated (Fig. 1B, *Repeat*). Repeats occurred at multiple lag distances, represented as negative values from repeat presentation to the original presentation, ranging from −2 (e.g. ABC<u>B</u>EF) to −5 (e.g. ABCDE<u>A</u>). A lag distance of −1 was not used due to the lack of an intervening item. Distribution of Repeats across all possible positions: Pos1, 0%; Pos2, 0%; Pos3, 7.5%; Pos4, 50%; Pos5, 27.5%; Pos6, 15%.

*Skips (forward lags)*. OutSeq images were considered Skips when presented too early in the sequence (e.g. AB<u>E</u>DEF; Fig. 1B, *Skip*). Skips occurred at all lag distances, represented as positive values from early presentation to original presentation, ranging from −1 (e.g. ABCD<u>F</u>F) to −4 (e.g. A<u>F</u>CDEF). Distribution of Skips across all possible positions: Pos1, 0%; Pos2, 15%; Pos3, 27.5%; Pos4, 50%; Pos5, 7.5%; Pos6, 0%.

### Sequence Memory Analysis

To evaluate whether participants demonstrated sequence memory, we compared the observed and expected frequencies of InSeq and OutSeq responses using *G*-tests (Sokal and Rohlf 1995). Responses to each item were sorted into a 2 × 2 matrix based on accuracy (correct/incorrect) and sequence condition (InSeq/OutSeq), as done previously (Allen et al., 2014; Jayachandran et al., 2019). For the sequence memory analysis, only “high memory” sequence sets were included. Responses to the first item of each sequence were excluded from analysis. Ordinal Transfers were excluded from this analysis because they were used to parse retrieval modes rather than measure performance. For comparison we evaluated sequence memory using d-prime, a measure of memory specificity derived from signal detection theory. Hits were defined as InSeq trials that were correctly identified as InSeq, misses were InSeq trials that were incorrectly identified as OutSeq, correct rejections were OutSeq probe trials that were correctly identified as OutSeq, and false alarms were defined as OutSeq probe trials that were incorrectly judged to be InSeq. The same conclusions were drawn with both approaches, thus, here we only report the results from the G-tests given its robustness to response biases as the current task has an overall greater number of InSeq responses.

Additionally, we examined overall sequence memory performance, using a summary statistic called the Sequence Memory Index (SMI; Equation 1). SMI normalizes the proportion of InSeq and OutSeq items across different conditions and represents sequence memory performance as a single value ranging from −1 to 1. A SMI of “1” represents perfect sequence performance (response time > 1 s for all InSeq items and < 1 s for all OutSeq items), and 0 represents chance performance. Note that an SMI score of −1 would indicate an incorrect response to every single item (response times < 1 s for all InSeq items and > 1 s for all OutSeq items). Negative SMIs rarely occurred, poor performance was typically captured by SMI scores close to zero. SMI was calculated for both low and high memory demand sequences and has been used in previous human studies to facilitate comparisons (Allen et al., 2014, 2015).

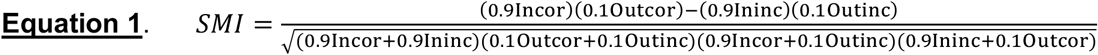

### Detailed Position and Lag Analysis

To evaluate strategies or mechanisms of sequence memory performance, a detailed analysis based on ordinal position and lag distance was performed for each OutSeq probe type (Ordinal Transfers, Skips, and Repeats). For Ordinal Transfers, we evaluated performance across positions (Pos2 thru Pos6). For Skips, we evaluated performance across *n*-forward lag distances (Fig. 3A; *n*-forward lags: −1, −2, −3, and −4). Smaller *n*-forward lags occurred more often because more combinations were available. For Repeats, we evaluated performance across the *n*-reverse lags (Fig. 3A; *n*-reverse lags: −2, −3, −4, and −5). Smaller *n*-reverse lags occurred more often because more combinations were available. We compared each position or lag distance performance (accuracy and SMI) using repeated-measures ANOVAs followed by one-sample t-tests. To account for the potential of a response bias (the assumption by participants of each item being in sequence) in the accuracy measures, we calculated an adjusted chance level as follows: (1) we calculated the observed response bias for holding the button for >1 sec on Pos 2 through Pos 6, irrespective of the InSeq or OutSeq status, which was 90.466%, and (2) we calculated the complement of that response bias and set that as the chance response for probe trials, which was 9.534%. To assess differences in variability amongst probe trial types trial specific performance was converted to a z-score across positions (Ordinal Transfers) and lags (Skips and Repeats) for each participant. Three mixed general linear models were performed with subject as a random effect and position or lag as a fixed effect. The resulting residuals were subsequently squared and averaged across positions or lags resulting in probe trial type specific residuals for each participant. We conducted a repeated measures ANOVA with Greenhouse-Geisser corrections and LSD posthoc testing to compare the averaged squared residuals across probe trial types (Ordinal Transfers, Skips and Repeats).

**Figure 2.**
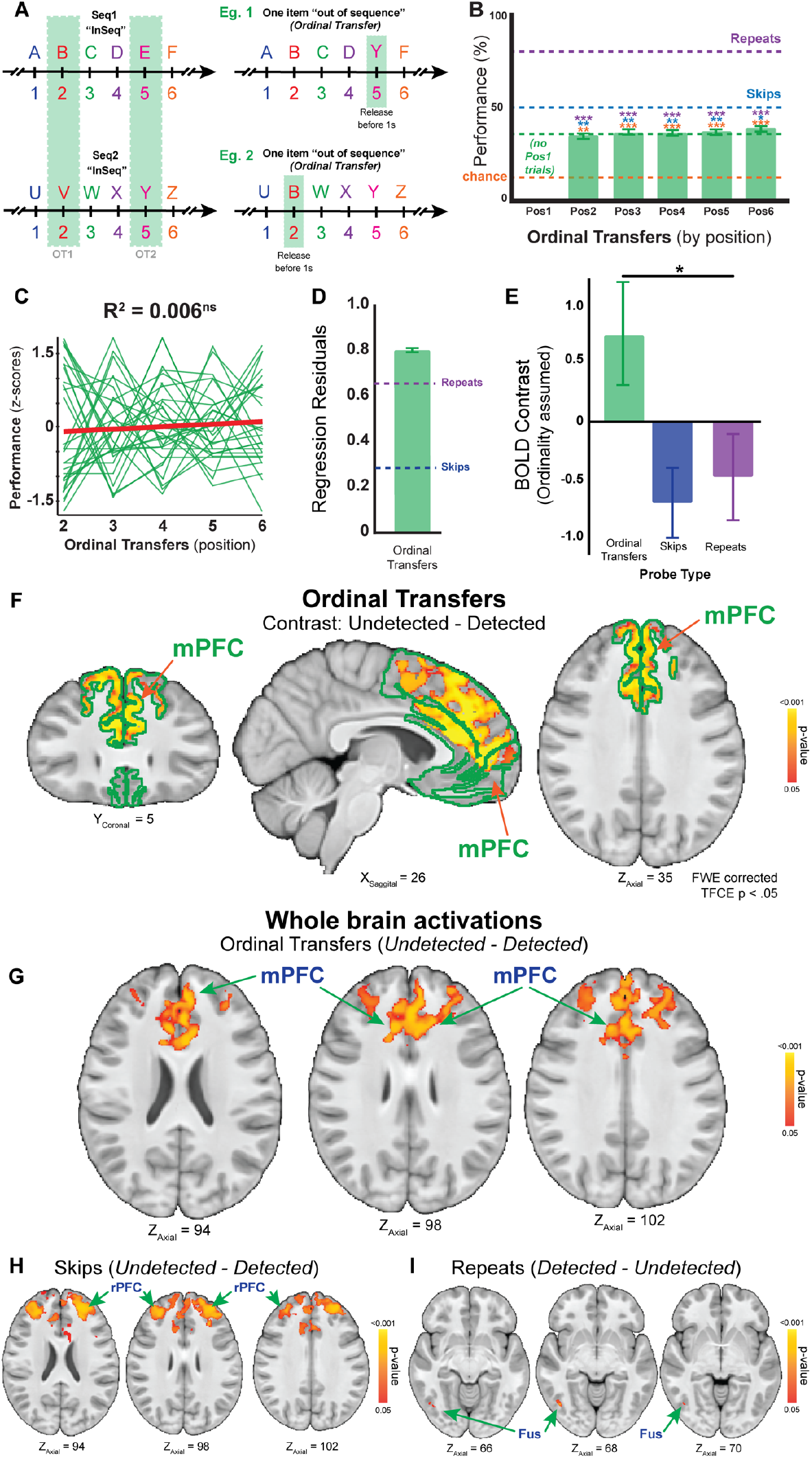
mPFC activations and ordinal retrievals. **A,** Ordinal retrieval modes were probed using Ordinal Transfer trials. Ordinal Transfers occurred when an item from one sequence (e.g., UVWXYZ) was transferred to another sequence (e.g., ABCDEF) while retaining in its original ordinal position (e.g. ABCD<u>Y</u>F). Theoretically, Ordinal Transfers are identified as InSeq when an ordinal association is used, and identified as OutSeq when a sequential item-item association is used. **B,** Ordinal Transfers occurred in every position in each sequence except the first. For each position (2 through 6), mean performance was significantly higher than chance (red dashed line; one minus the calculated response bias), and significantly lower than the mean performance on Repeats (purple dashed line) and Skips (blue dashed line). Performance did not significantly differ among positions. **C,** the normalized performance slope did not significantly differ across positions, and variability, measured by the averaged squared residuals per participant of the linear regression, was high overall. High performance variability suggests conflicting strategies (Repeats and Skips variability levels plotted for comparison in **D**). For the subsequent BOLD analysis, we focused on contrasts assuming participants were using an ordinal retrieval mode. When an ordinal retrieval mode is engaged, participants would identify Ordinal Transfers as InSeq, but would identify Repeats and Skips as OutSeq. **E**, a BOLD fMRI mPFC ROI analysis showed that the Ordinal Transfers Undetected – Detected contrast activation was significantly higher compared with Repeats and Skips (Detected – Undetected Contrasts). **F,** a voxel-wise BOLD fMRI analysis using mPFC as a mask (outlined in green) resulted in significant activity in a large area of the mPFC. To investigate whether there was a difference between these probe trials, we compared Ordinal Transfers Undetected – Detected, Repeats Detected – Undetected, and Skips Detected – Undetected trials. **G,** mPFC activation was also evident in whole brain analysis for Ordinal Transfers Undetected – Detected. **H,** additionally, we found activation when comparing Skips Undetected – Detected in the rostrolateral prefrontal cortex. No significant activation was found when comparing Repeats Undetected – Detected (not shown). ***I***, when comparing Repeats Detected – Undetected we found activation in the right fusiform gyrus. No significant activation was found when comparing Skips Detected – Undetected (not shown). Notably, mPFC activation did not simply reflect inaccuracy on OutSeq trials. Symbol(s): ns, not significant; *, p < 0.05, **, p < 0.01, *** p < 0.001.

**Figure 3.**
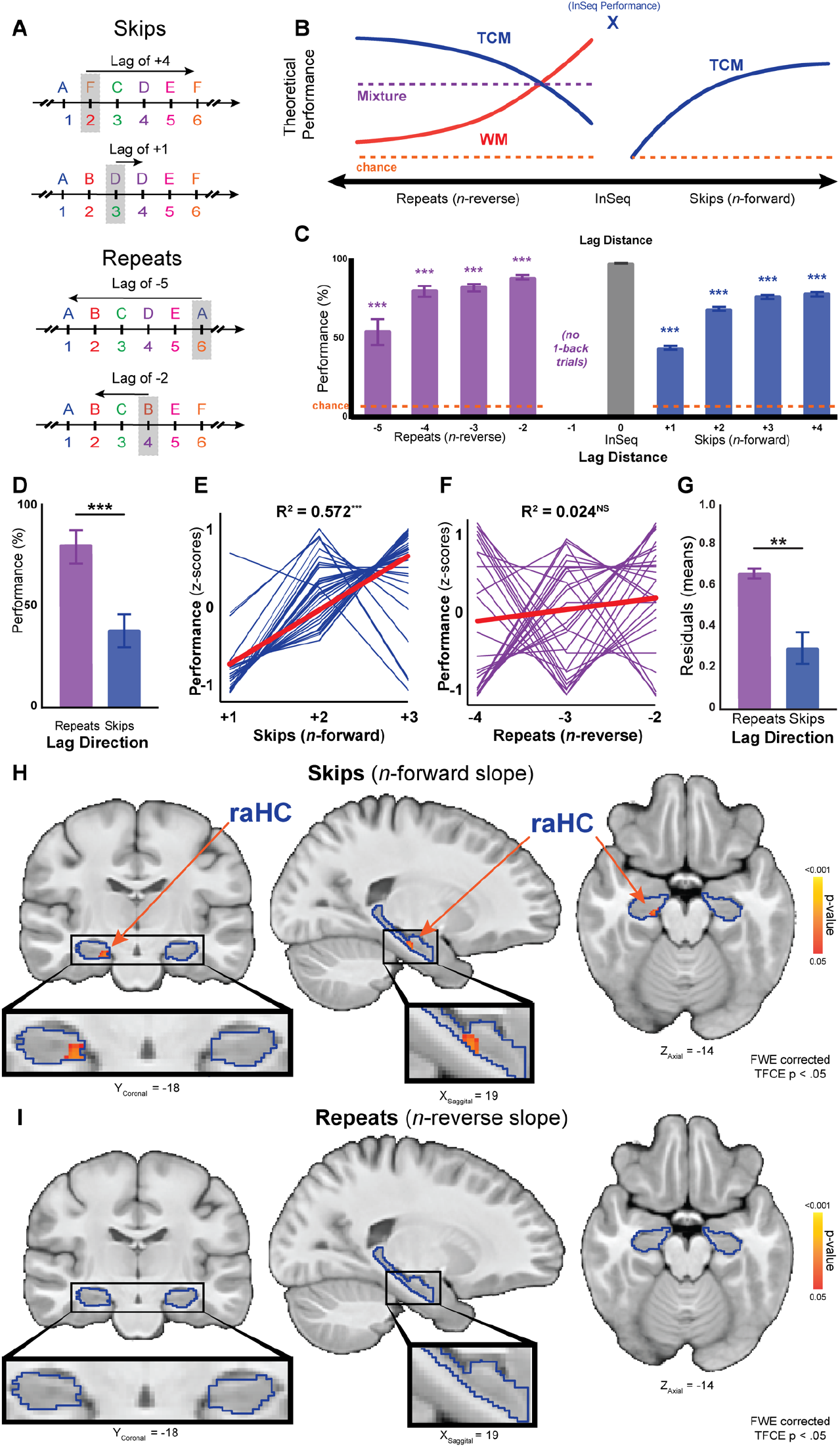
BOLD fMRI analysis indicates that the right anterior hippocampus (raHC) tracks temporal context across forward lags. **A,** Skips and Repeats could occur at different lags away from their InSeq position. Skip lags ranged from −1 (e.g. ABCD<u>F</u>F) to −4 (e.g. A<u>F</u>CDEF). Repeat lags ranged from −2 (e.g. ABC<u>B</u>EF) to −5 (e.g. ABCDE<u>A</u>). ***B,*** theoretical predictions based on temporal context memory (TCM) and working memory (WM) retrieval modes for different lag distances on Repeats (e.g., AB<u>A</u>) and Skips (e.g., AB<u>D</u>). The TCM (blue lines) predicts that item retrieval in sequences is graded by lag distance (for both Repeats and Skips). According to the TCM, OutSeq probe trials with small lags would be more difficult to detect than those with larger lags reflecting the fact that they were more temporally contiguous in the original sequence and likely to be judged InSeq. Conversely, the WM (red line) predicts that items with small lags would be easier to detect than those with larger lags, but WM can only contribute to performance on Repeats. A mixture of TCM and WM modes is shown by the dashed purple line. **C,** the mean performance accuracy for all participants on Repeats and Skips at different lags. **D,** we predicted and found that overall performance on Repeats would be higher than for Skips because TCM and WM retrieval modes complement each other. **E,** we also predicted and found that performance would show a positive slope on Skips defining of TCM retrieval modes, **F,** and would be flat on Repeats reflecting the use of a mixture of TCM and WM retrieval modes. Specifically, a linear regression in Skips performance positively increased across forward lags, but did not have a significant slope on Repeats. **G,** finally, we predicted and found significantly higher variability, measured by the averaged squared residuals per participant of the linear regression, in Repeats compared with Skips, which is thought to reflect the integration of variance from using both TCM and WM retrieval modes. Symbol(s): ns, not significant; *, p < 0.05, **, p < 0.01, *** p < 0.001. **H,** BOLD fMRI analysis of Skips with a linear contrast of − 1, 0, 1 for lags of −1, −2, −3 respectively using the bilateral medial temporal lobe as a revealed significant activation in the raHC (right anterior Hippocampus). ***I***, BOLD fMRI analysis of Repeats with a linear contrast of 1, 0, −1 for lags of −4, −3, −2 respectively using bilateral medial temporal lobe as a mask showed no significantly activated clusters.

### Neuroimaging Acquisition

Neuroimaging data were collected on a General Electric Discovery MR750 3.0T scanner using a 32-channel head coil at the University of Miami Neuroimaging Facility. Structural T1-weighted images were collected (186 slices, flip angle = 12°; TE = 3.68 ms, TR = 9.184 ms, TI = 650 ms, matrix = 256 × 256 mm; FOV = 256 mm, slice thickness = 1.0 mm). Whole-brain T2*-weighted, blood oxygen level dependent (BOLD) echo-planar imaging data (42 slices, interleaved, bottom-up; flip angle = 75°; TE = 25 ms; TR = 2000 ms; matrix = 96×96 mm; FOV = 240 mm, slice thickness = 3.0 mm, voxel size = 2.5 mm^2^) were collected during the task.

### Neuroimaging Preprocessing

Preprocessing was performed using a pipeline developed in Nipype v0.1 (Gorgolewski et al., 2011), wrapping tools from Analysis of Functional Neuroimages (AFNI v16.3.18; Cox, 1996), FSL (v5.0.10), FreeSurfer (v5.1.0; Dale et al., 1999), Advanced Normalization Tools (v2.2.0; Avants et al., 2008), and Artifact Detection Tools (ART; www.nitrc.org). Following DICOM conversion, cortical surface reconstruction and cortical/subcortical segmentation was performed on all T1-weighted structural scans. Functional data were first ‘despiked’ to remove and replace intensity outliers in the functional time series. The data were simultaneously slice-time and motion corrected (Roche, 2011). An affine transformation matrix was calculated during coregistration of the mean of each participant’s functional scans to their structural scan using Freesurfer’s boundary-based registration algorithm (BBregister). Brain masks were created by binarizing the aparc−aseg file and dilating by one voxel. The resulting binary masks were subsequently coregistered to the functional data by applying the inverse of the affine coregistration matrix. Motion and intensity outliers were then identified using the rapid art artifact detection tool as implemented in nipype. Time-points at which intensity either exceeded three standard deviations or composite frame-wise displacement was greater than 1 mm were flagged as outliers to serve as subsequent regressors of no interest in the first-level general linear models. Finally, functional data were spatially filtered with a 5 mm FWHM maximum Gaussian kernel using the *SUSAN* algorithm (FSL).

### Neuroimaging Normalization

A study-specific template was generated using Advanced Normalization Tools (ANTs). Each structural scan was skull-stripped by multiplying the T1-weighted structural scan by the binarized and dilated aparc−aseg file in structural space. Each skull-stripped brain was then rigid-body transformed (no scaling or shearing) to Montreal Neurological Institute space using FSL’s *FLIRT* algorithm. This first pass was used to minimize large spatial shifts between participants and generate a template close to a commonly used reference. Following visual inspection, a study template was created using the buildtemplateparallel.sh script from ANTs. After template generation, each participant’s skull-stripped brain was normalized using non-linear symmetric diffeomorphic mapping implemented by ANTs. The resulting warps were applied to contrast parameter estimates following fixed-effects modeling for subsequent group-level tests.

### Neuroimaging Analysis

Functional data were analyzed according to a general linear model approach using FMRIB’s Software Library (www.fmrib.ox.ac.uk/fsl). Two separate univariate general linear models were used for first-level analyses: (1) a performance (correct/incorrect) model; and (2) a lag model.

All first-level models included event and nuisance regressors. The performance analysis contained eight event regressors of interest: correct and incorrect InSeq, Repeats, Skips, and Ordinal Transfers. Event regressors were convolved with FSL’s double gamma hemodynamic response function with an onset beginning at the first stimulus item of a sequence and a duration equal to the length of time required to evaluate all the items in that sequence (mean = 9.917 s per subject). The lag analysis comprised six event regressors of interest: Repeats at different *n*-reverse lags (*n*-reverse lags: −2, −3, and −4), and Skips at different *n-*forward lags (*n*-forward lags: −1, −2, and −3). InSeq and Ordinal Transfers irrespective of behavioral performance were also included. We evaluated linear changes in activation across lags with the following contrast weights: −1, 0, 1 and 1, 0, −1 for Repeats (*n*-reverse lags: −2, −3, and −4) and Skips (*n*-forward lags: −1, −2, and −3). We did not include a lag of −5 or −4, as the numbers of trials with these lags per participant was not sufficient (1 trial). Regressors for baseline or low memory sequences were also included in both models. Nuisance regressors included motion parameters (x, y, z translations; pitch, roll, yaw rotations), first and second derivatives of the motion parameters, normalized motion, first, second, and third order Lagrange polynomials, as well as each outlier time-point that exceeded the artifact detection thresholds identified during preprocessing.

Following the first-level analysis, a fixed effects analysis across experimental runs was performed for each participant for the respective contrasts of interest (e.g., performance analysis: correct versus incorrect probe trials; lag analysis: linear contrasts). Contrast parameter estimates from the fixed effects analysis were normalized to the study-specific template and group-level analyses were performed using FSL’s *randomise* threshold-free cluster enhancement (tfce) one sample t-test. To test *a priori* hypotheses with respect to the functional contributions of the mPFC and HC during sequence memory retrieval, we constrained our voxel-wise analyses at the group level to the bilateral mPFC and medial temporal lobe using masks of these regions. Whole-brain exploratory analyses were used to follow up our anatomically directed tests. In the performance analysis, some participants were missing key events of interest (e.g., correct Ordinal Transfers or incorrect Repeat probes). These participants were not included in the relevant analyses, reducing the sample size for the correct versus incorrect Ordinal Transfer contrast to n = 26, and for the correct versus incorrect repeat probe trial contrast to n = 31.

To directly examine the relation between sequence memory retrieval mode and regional contribution, as a post hoc follow up to our voxelwise analyses, we used an anatomical region of interest analysis to perform a 2×2 repeated measures factorial ANOVA with brain region (right anterior HC vs. mPFC) and strategy (ordinal retrieval mode [Ordinal Transfer incorrect > Ordinal Transfer correct] vs. temporal context retrieval mode [TCM; Positive Linear Skip contrast]) as the within subjects factors and participants as the repeated measure. Anterior HC region of interest was manually drawn in coronal slices. The anterior most boundary of the hippocampus was defined by the white matter separating the hippocampus from the amygdala, the lateral and medial boundaries were defined by the cerebral spinal fluid of the lateral ventricle, the inferior boundary was white matter of the parahippocampal gyrus, while the superior boundary was the wavelike contour of the pes digitations/alveus/horizontal line connecting the middle of the medial border of the lateral ventricle to the surface of the uncus. The anterior HC was delineated from the posterior HC by the presence of a small anatomical protrusion of medial HC into the lateral ventricle that was absent in the posterior HC – uncal apex. The mPFC region of interest was created by binarizing the medial orbitofrontal, superior frontal, rostral anterior cingulate, and the caudal anterior cingulate Freesurfer labels.

## Results

### Sequence Memory

We first looked at overall sequence memory in the task by comparing performance on low and high memory demand sequence sets. We calculated an SMI separately for the low and high memory demand sequences (Fig. 1A). As expected, participants performed significantly better than chance (SMI = 0) on both low memory sequences (SMI_low_: 0.751 ± 0.173; SMI_low vs. chance_: t_(33)_ = 37.065, p = 3.487 × 10^−23^) and high memory sequences (SMI_high_: 0.582 ± 0.170; SMI_low vs. chance_: t_(33)_ = 19.690, p = 5.391 × 10^−20^), but performance was significantly better with low memory sequences (SMI_low vs. high_: t_(33)_ = −4.839, p = 2.958 × 10^−5^). High memory sequences were used for all subsequent analyses because they optimally taxed the different sequence retrieval modes. Importantly, there was no testing order effect (1^st^ or 2^nd^ experimental block) on sequence memory (SMI_1st block_ = 0.571 ± 0.190, SMI_2nd block_ = 0.594 ± 0.152; SMI_1st vs. 2nd_: t_(32)_ = −0.386, p = 0.702), nor did we observe any sex differences (SMI_female_ = 0.566 ± 0.197, SMI_male_ = 0.602 ± 0.133; SMI_female vs. male_ = t_(32)_ = −0.612, p = 0.545). Thus, we pooled these groups.

Next, we evaluated overall performance across each of the three memory probes individually using percent correct for ease of interpretation. As in previously published studies, we found that Ordinal Transfers (Accuracy = 34.927 ± 4.120%) were the most difficult, followed by Skips (Accuracy = 49.205 ± 2.686%), with Repeats being the easiest (Accuracy = 78.584 ± 2.853%). There was a significant difference in performance across probe types (F_(2,66)_ = 85.714, P = 3.433 × 10^−14^) suggesting differences in the available retrieval modes driven by the different conditions. Post hoc pairwise comparisons showed that the performance on Repeats was significantly higher compared to Skips (mean difference = 0.294 ± 0.020, p = 8.137 × 10^−16^) and Ordinal Transfers (mean difference = 0.437 ± 0.039, p = 1.078 × 10^−12^) and performance on Skips was significantly higher compared to Ordinal Transfers (mean difference = 0.143 ± 0.039, p = 0.001). The same pattern of results was evident when using SMI which controls for idiosyncratic response patterns. Importantly, participants performed each of the three probe types significantly better than chance (Ordinal Transfers: t_(33)_ = 6.164, p = 5.951 × 10^−^ 7, Repeats: t_(33)_ = 24.201, p = 1.403 × 10^−22^; Skips: t_(33)_ = 14.771, p = 4.203 × 10^−16^). For all probe types, learning was rapid (asymptotic within a few trials), and performance was steady throughout the duration of the experiment (Fig. 1C). Thus, performance in the task and the accompanying brain imaging data primarily reflect distinct memory retrieval modes driven by specific probe conditions.

### Ordinal Memory Retrievals

An ordinal retrieval mode is known to contribute to memory for sequences of events which we tested for here using Ordinal Transfer probe trials (e.g., A**<u>2</u>**CDEF, Fig. 2A). If participants *exclusively* used an ordinal retrieval mode on these trials (e.g., A goes in the 1^st^ position, B goes in the 2^nd^ position, etc.), then these probes would be remembered as being InSeq (since “2” is in the same position as B). An ordinal retrieval mode would be expected to drive performance on these trials to very low levels, possibly to chance, if no other retrieval mode was engaged. Moreover, if an ordinal retrieval mode was used, we would not expect differences in accuracy as a function of transfer position. Conversely, if participants relied on another process, such as a temporal context, then Ordinal Transfers would be easily identified as OutSeq (since “2” does not follow A, and in fact it doesn’t go with any of the items in that sequence). Non-ordinal retrieval modes would drive performance to very high levels on Ordinal Transfer probe trials, probably to the same levels as at Repeats (as a good empirical benchmark for asymptotic performance).

We found Ordinal Transfers were performed significantly better than chance across all positions (Fig. 2B, red stars; Pos2: 0.314 ± 0.357; Pos2_vs. chance_: t_(33)_ = 3.566, p = 0.001; Pos3: 0.338 ± 0.313; Pos3_vs. chance_: t_(33)_ = 4.512, p = 7.712 × 10^−5^; Pos4: 0.347 ± 0.258; Pos4_vs. chance_: t_(33)_ = 5.670, p = 2.549 × 10^−6^; Pos5: 0.354 ± 0.283; Pos5_vs. chance_: t_(33)_ = 5.334, p = 6.892 × 10^−6^; Pos6: 0.373 ± 0.336; Pos6_vs. chance_: t_(33)_ = 4.811, p = 3.212 × 10^−5^) suggesting participants utilized non-ordinal retrieval modes for these trials. However, performance was much lower than Repeats (Fig. 2B, purple stars; Pos2_vs. Repeats_: t_(33)_ = − 7.711, p = 7.019 × 10^−9^; Pos3_vs. Repeats_: t_(33)_ = −8.338, p = 1.224 × 10^−9^; Pos4_vs. Repeats_: t_(33)_ = −9.917, p = 1.996 × 10^−11^; Pos5_vs. Repeats_: t_(33)_ = −8.919, p = 2.621 × 10^−10^; Pos6_vs. Repeats_: t_(33)_ = −7.175, p = 3.185 × 10^−8^) and lower than Skips (Fig. 2B, blue stars; Pos2_vs. Skips_: t_(33)_ = −2.912, p = 6.390 × 10^−3^; Pos3_vs. Skips_: t_(33)_ = − 2.870, p = 7.104 × 10^−3^; Pos4_vs. Skips_: t_(33)_ = −3.284, p = 2.426 × 10^−3^; Pos5_vs. Skips_: t_(33)_ = −2.854, p = 7.400 × 10^−3^; Pos6_vs. Skips_: t_(33)_ = −2.075, p = 4.589 × 10^−2^) suggesting a heavy reliance on an ordinal retrieval mode. We found no significant difference in performance across positions (Fig. 2B; F_(4, 132)_ = 0.331, p = 0.786). The observed behavioral performance suggests that participants were disproportionally using an ordinal retrieval mode (e.g., lower performance when compared to Repeats and Skips) in combination with other non-ordinal retrieval modes (i.e., still at performance better than chance) to a lesser extent. A prediction from the these conflicting retrieval modes is that performance variability would be high between participants and across positions. In line with this hypothesis we observed higher Ordinal Transfer variability when compared to Repeats, and much higher when compared to Skips (Fig. 2C-D; M_residual_ = 0.795 ± 0.074). A repeated-measure ANOVA with Greenhouse-Geisser correction showed significant main effect among regression residuals of Ordinal Transfers, Skips and Repeats (F_(2, 54)_ = 23.866, p = 1.843 × 10^−8^). Regression residuals of Ordinal Transfers were significantly higher compared to Skips (mean difference = 0.530 ± 0.089, p = 2.523 ×10^−6^) and Repeats (mean difference = 0.158 ± 0.028, p = 5.627×10^−6^). Regression residuals of Repeats were significantly higher compared to Skips (mean difference = 0.373 ± 0.099, p = 0.001).

### mPFC Activations and Ordinal Retrievals

We hypothesized that using an ordinal retrieval mode would be related to activations in the mPFC (e.g., Hsieh and Ranganath, 2015). Algorithmically, this would be evident when contrasting undetected (i.e., using an ordinal retrieval mode) versus detected (i.e., using a non-ordinal retrieval mode) during Ordinal Transfer trials. We also reasoned that the use of an ordinal retrieval mode would facilitate the detection of Skips and Repeats (because of the ordinal position mismatch), thus, when assuming an ordinal retrieval mode for Skips and Repeats we evaluated activations contrasting detected versus undetected trials. We used an anatomical region of interest (ROI) analysis to evaluate mPFC activity across the three probe trial types. In support of our hypothesis the mPFC was significantly more active during identification of Ordinal Transfers relative to both Skips and Repeats for contrasts that assumed ordinal retrieval modes (Fig. 2E; F_(2, 56)_ = 4.038, p = 0.034). Post hoc tests (LSD) further supported the conclusion that the mPFC was most active in the comparison of undetected versus detected Ordinal Transfers (M = 0.708 ± 1.998) relative to detected versus undetected Skips (M = −0.649 ± 1.445, p = 0.051) and Repeats (M = −0.431 ± 1.728, p = 0.005). No significant difference was identified in mPFC activation between detected versus undetected Skips and Repeats (mean difference = 0.061 ± 0.437, p = 0.890). As a follow-up and to explore contributions of distinct regions within our anatomical mPFC ROI we evaluated the same contrasts at the voxel-wise level. We observed activations throughout the mPFC bilaterally (prelimbic cortex, anterior cingulate cortex, medial superior frontal gyrus) following the comparison of undetected versus detected Ordinal Transfers (Fig. 2F; FWE-tfce p < 0.05), while no mPFC clusters survived corrections for multiple comparisons when comparing detected versus undetected Skips (data not shown) and Repeats (Fig. 2I; FWE-tfce p > 0.05).

One potential confound to this interpretation is that mPFC activations might reflect relative performance levels (incorrect > correct trials) rather than the utilization of ordinal retrieval modes per se. If true, we would expect similar mPFC activation clusters when examining incorrect compared to correct Skips and Repeats if performance was the main contributor to activation. When evaluating performance related activations for Repeats and Skips, undetected (i.e., incorrect) compared to detected (i.e., correct) Skips exhibited activations predominantly in rostrolateral prefrontal cortex (Fig. 2G) while a cluster for detected greater than undetected Repeats was identified in the right fusiform gyrus (Fig. 2H). These results further support the view that the mPFC contributes to ordinal retrieval modes, and do not generally reflect performance accuracy. Notably, the observed activations in the rostrolateral prefrontal cortex during undetected Skips, and the fusiform gyrus for detected Repeats may reflect prospective memory (Umeda et al., 2011; Volle et al., 2011; Benoit et al., 2012) and object processing (Grill-Spector et al., 1998), respectively.

### Sequence memory as a function of lag direction and distance

Memory for sequences of events at different lags can be supported by a variety of cognitive processes including working memory (WM) and temporal context memory (TCM) retrieval modes. The use of WM and TCM retrieval modes predict different patterns in behavioral performance across n-forward and n-reverse lags (see Fig. 3B; also see Jayachandran et al., 2019), which can be exploited here to test the use of different retrieval modes (e.g., WM versus TCM).

*Skips (forward lags).* OutSeq probe trials that skipped ahead in the sequence (n-forward, e.g., the “D” in ABDDEF) afford the opportunity to evaluate predictions of the use of a TCM retrieval mode during rapid sequence memory decisions. Specifically, successful performance on Skips relies on a TCM retrieval mode as it requires participants to have precise expectations for the subsequent items in the sequence. Accordingly, the likelihood of a memory retrieval is highest for the very next item in the forward direction (lag distance = −1) and drops off (in a graded fashion) for more distal items (from −2 to −4). In the context of this task, the use of a TCM retrieval mode predicts the inverse in performance compared with free recall tasks because of interference. Specifically, performance on Skip OutSeq probe trials should be most difficult to detect for the shortest forward lag distance (−1) and improve at longer distances (−2, −3, or −4; Fig. 3B, right side) precisely because proximal items in a sequence (i.e., Skips with a short forward lag) are more likely to be retrieved (Howard and Kahana 2002; Kragel et al. 2015) and then falsely match up with an out of sequence probe image (thus be judged as InSeq; undetected).

We tested this prediction by examining performance across n-forward lag distances. First, all forward lags were performed better than chance (Fig. 3C, blue bars; −1: 0.420 ± 0.178; −1_vs. chance_: t_(33)_ = 10.609, p = 3.608 × 10^−12^; −2: 0.665 ± 0.156; −2_vs. chance_: t_(33)_ = 21.299, p = 7.417 × 10^−21^; −3: 0.747 ± 0.248; −3_vs. chance_: t_(33)_ = 15.343, p = 1.402 × 10^−16^; −4: 0.765± 0.431; −4_vs. chance_: t_(33)_ = 9.066, p = 1.780 × 10^−10^), suggesting that the use of a TCM retrieval mode does not completely interfere with OutSeq detection at the different forward lag distances. Second, subjects exhibited graded performance improvement as the skip distance increased (F_(3, 99)_ = 13.790, p = 1.379 × 10^−4^). We found that the identification of Skips with a −1 lag were the most difficult to detect compared with all other forward lags (*post hoc* LSD: −2: mean difference = 0.245 ± 0.029, p = 1.218 × 10^−9^, −3: mean difference = 0.327 ± 0.049, p = 1.248 × 10^−7^, −4: mean difference = 0.345 ± 0.079, p = 1.186 × 10^−4^). These results are consistent with TCM retrieval mode predictions (Fig. 3B, 3C). Detection of Skips with a −2 lag was significantly lower than Skips with a −3 lag (*post hoc* LSD: mean difference = 0.082 ± 0.038, p = 0.039). No significant difference in detection was observed, however, when comparing Skips with lags of −2 and −4 (*post hoc* LSD: mean difference = 0.100 ± 0.074, p = 0.187), or −3 and −4 (*post hoc* LSD: mean difference = 0.018 ± 0.074, p = 0.814), suggesting that performance approached asymptote. These behavioral results suggest that the participants’ sequence memory was driven by the use of a TCM retrieval mode on Skips.

*Repeats (reverse lags).* OutSeq trials that repeated an item from earlier in the sequence (*n*-reverse; e.g., the second “A” in AB<u>A</u>DEF) can be solved by either the use of a TCM retrieval mode, a WM retrieval mode, or a mixture of the two (Fig. 3B, left side). Notably, the pattern of OutSeq detections across *n*-reverse lag distances should differentiate the two competing processes (Jayachandran et al., 2019). For Repeats, a TCM retrieval mode predicts that short backward lag distances (e.g., −2) would be the most difficult to detect due to heightened interference, while further *n*-reverse lag distances (e.g. −5) would be readily detectable as OutSeq. By contrast, a WM retrieval mode predicts the opposite pattern because the most recently experienced items would be the most accessible and therefore easiest to detect.

We found that performance was better than chance for all *n*-reverse lag distances (Fig. 3C, purple bars, −2: 0.859 ± 0.116; −2_vs. chance_: t_(33)_ = 38.333, p = 6.058 × 10^−29^; −3: 0.805 ± 0.171; −3_vs. chance_: t_(33)_ = 24.138, p = 1.522 × 10^−22^; −4: 0.782 ± 0.233; −4_vs. chance_: t_(33)_ = 17.204, p = 4.821 × 10^−18^; −5: 0.529 ±0.507; −5_vs. chance_: t_(33)_ = 4.996, p = 1.865 × 10^−5^). As the *n*-reverse lag distance increased, detection as OutSeq decreased (F_(3, 99)_ = 9.760, p = 0.001; Fig. 3C, purple bars), suggestive of the dominant use of a WM retrieval mode. Items with an *n*-reverse lag closest to their original position (e.g., ABC<u>B</u>EF; lag = −2) were easiest to detect as OutSeq compared with all other *n*-reverse lag positions (*post hoc* LSD lag −2 compared with: lag −3: mean difference = 0.055 ± 0.022, p = 0.020; lag −4: mean difference = 0.077 ± 0.033, p = 0.025; or lag −5: mean difference = 0.330 ± 0.084, p = 4.262 × 10^−4^). By contrast, items with an *n*-reverse lag farthest from their original position (e.g. ABCD<u>E</u>A; lag = −5) were the most difficult to detect as OutSeq compared with all other *n*-reverse lag positions (*post hoc* LSD lag −5 compared with: lag −4: mean difference = 0.253 ± 0.091, p = 0.009; lag −3: mean difference = 0.275 ± 0.091, p = 0.005; or lag −2: mean difference = 0.330 ± 0.084, p = 4.262 × 10^−4^). No significant differences in detection between −3 and −4 *n*-reverse lags were observed (*post hoc* LSD: mean difference = 0.022 ± 0.035, p = 0.532). Taken together, performance at the *n*-reverse lag extremes (−2 and −5) supports the notion that the use of a WM retrieval mode plays an important role in identifying Repeats. The absence of a graded performance across *n*-reverse lags of −3 and −4 supports the idea that a combination of cognitive processes is being used, but overall these analyses do not support the use of a TCM retrieval mode as an isolated process driving the identification of Repeats in this task.

### TCM retrievals for Skips, but a mixture of retrieval modes for Repeats

Differences in the overall ability to detect OutSeq probe trials, and the residuals from a lag-based linear regression model, helped to further elucidate the contributions of either a TCM retrieval mode, a WM retrieval mode, or their combination. First, we predicted that the ability to detect OutSeq probes would be greater for Repeats than Skips because TCM and WM retrieval modes can both contribute to the evaluation of Repeats but not Skips (Fig. 3B, left, purple dashed line). Second, we predicted that a linear regression of Skip detections would positively increase across *n*-forward lags, whereas a similar analysis across *n*-reverse lags for Repeats would be essentially flat. Third, we predicted that the residuals from the linear regressions would be, highest on Repeats compared with Skips, reflecting the use of multiple retrieval modes, whereas Skips involve a single mode (i.e., TCM-based retrieval). To test these predictions, we compared the overall ability to detect the two probe trial types, calculated a linear regression based on the z-scores (accounting for individual baseline detection levels) across different lag positions, and averaged the squared residuals as a measure of detection variability across lags. Participants were significantly better at detecting Repeats (M_accuracy_ = 0.786 ± 0.166) than Skips (M_accuracy_ = 0.492 ± 0.157; t_(33)_ = 14.435, p = 8.137 × 10^−16^; Fig. 3D).

Consistent with our hypothesis, the linear regression in Skips positively increased across forward lags accounting for a large effect on performance (Fig. 3E; R^2^ = 0.572, β = 0.757, p = 3.711 × 10^−20^), whereas a linear regression across reverse lags for Repeats had no significant slope (Fig. 3F; R^2^ = 0.024, β = 0.154, p = 0.139). When quantifying variability, Repeats (M_residual_ = 0.593 ± 0.243) were significantly more variable compared with Skips (M_residual_ = 0.285 ± 0.478) across different lag positions (Fig. 3G; t_(30)_ = 3.051, p = 0.004). The patterns in OutSeq detection and variability further support the conclusion that multiple retrieval modes likely contribute to identifying Repeats as OutSeq, while Skip detection is mediated more exclusively by a TCM retrieval mode.

### HC activations and TCM retrievals

Converging evidence indicates that the medial temporal lobe, specifically the HC formation, plays a disproportionate role in the use of a TCM retrieval mode in the brain (Manns et al., 2007; Hsieh et al., 2014; Bladon et al., 2019). To test whether regions of the medial temporal lobe contribute to a TCM retrieval mode during Skip and Repeat probe trials, we looked for linear changes in activation across the different lag distances. We observed a significant activation cluster in the right anterior HC following small volume corrections (bilateral medial temporal lobe, FWE-tfce p < 0.05) that increased its activation across n-forward lags (Fig. 3H). The statistical significance of similar patterns of activation across n-reverse lags did not survive corrections for multiple comparisons (Fig. 3I). These results suggest that the right anterior HC contributes to a TCM retrieval mode during Skips.

### Brain region by retrieval mode interaction effect

After finding individual evidence supporting the mPFC contributes to sequence memory by the use of an ordinal retrieval mode and the right anterior HC by a TCM retrieval mode, we wanted to directly compare the mode-related activations (ordinal vs. TCM) across the regions that exhibited voxelwise activations (right anterior HC vs. mPFC). A repeated-measures factorial ANOVA was conducted with brain region (anatomically defined right anterior HC vs. mPFC) and strategy (ordinal retrieval vs. TCM retrieval modes) as within-subjects factors, and BOLD activations as the dependent measure. We observed greater activations in the mPFC in relation to the use of an ordinal retrieval relative to a TCM retrieval mode, and the opposite pattern in the right anterior HC, evidenced by a significant interaction effect (Fig. 4A. F_(1,31)_ = 4.782, p = 0.036).

**Figure 4.**
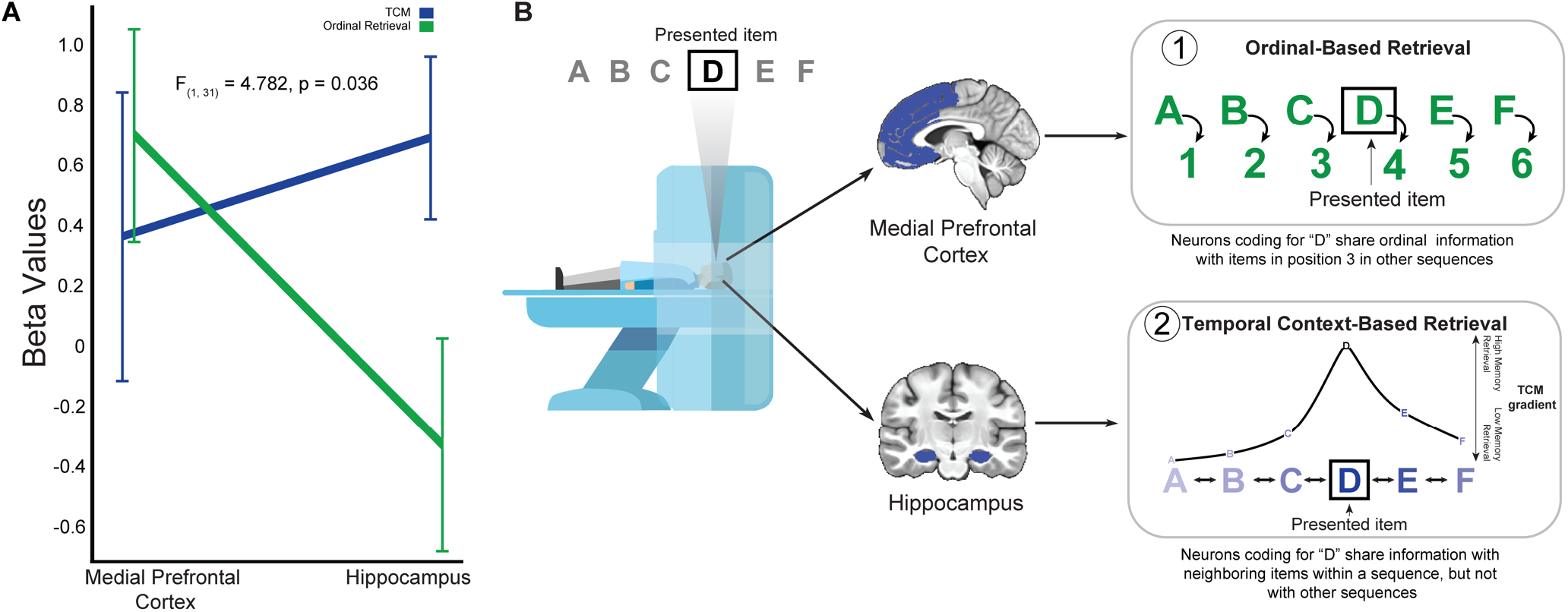
mPFC and HC retrieval mode interactions. **A,** there was a significant interaction effect between mode and brain region (F(1,31) = 4.782, p = 0.036), where we observed greater activations in the mPFC for ordinal retrievals relative to TCM retrievals, and the opposite pattern in the right anterior HC. **B,** based on the results of the current experiment, **1**) the mPFC engages in ordinal-based retrieval calling on associations between items and their ordinal position (which could be represented elsewhere) and **2)** the HC engages in temporal context-based retrieval.

## Discussion

The current study used different out of sequence probe trials during a memory task to test the concurrent use of distinct retrieval modes, adding to a growing literature on sequence memory as a fundamental component of episodic memory (Tulving, 1984; 2002; Allen and Fortin, 2013; Howard and Eichenbaum, 2013; Eichenbaum, 2017). Complementary behavioral results across the different out of sequence probe trials in the current task support the conclusion that memory for sequences of events is supported by both ordinal and temporal context retrieval modes. The BOLD fMRI evidence showed that mPFC activations reflect the use of an ordinal retrieval mode, while activations in the HC reflected a temporal context memory retrieval mode, thus dissociating the neurobiological substrates of two distinct processes contributing to sequence memory. While considerable evidence indicates that the mPFC and HC are involved in sequence memory, the current study provides new results that the mPFC and HC are concurrently engaged in different retrieval modes in support of remembering *when* an event occurred. This evidence provides an important baseline for further investigation of how sequence memory is impaired in typical aging and diseases such as Alzheimer’s Disease on the neurobiological level, especially since evidence suggests the relative dependence on an ordinal retrieval mode increases while the use of a temporal context retrieval mode declines with age (Bastin and Van der Linden, 2010; Allen et al., 2015).

### Ordinal retrieval modes in mPFC

Our results suggest that the positional information associated with retrieving memories in sequence is represented in the mPFC (e.g., Hsieh and Ranganath, 2015), and/or that mPFC activations help engage these representations elsewhere such as in HC neurons (e.g., Allen et al., 2016). Ordinal Transfers were detected less often than both Repeats and Skips (Fig 2B, see also Allen et al., 2014, 2015), but better than chance, suggestive of the use of multiple retrieval mode perhaps concurrent ordinal and temporal context retrieval modes. These results are not totally surprising because mPFC has been shown to be generally important to temporal order memory (Milner et al., 1985; Shimamura et al., 1990; DeVito and Eichenbaum, 2011; Hsieh et al., 2015) and other semantic representations (Preston and Eichenbaum, 2013; Hyman et al., 2012). Our study shows that when an ordinal retrieval mode is strongly engaged, it interferes with the ability to detect Ordinal Transfers, supported by the fact that Transfers went undetected at a very high rate, and mPFC activation was similarly high in such a way that could not be attributed to undetected probes more globally.

Ordinal retrieval modes have been demonstrated in rats (Allen et al., 2014), monkeys (Orlov et al., 2000; 2006) and humans (Allen et al., 2015; Hsieh et al., 2015). In fact, monkeys natural and dominant strategy is to categorize items within a sequence by their ordinal position, a strategy that occurred first in Orlov et al. (2000), and only later in trials did monkeys employ other strategies such as sequential associations for adjacent items and working memory (Orlov et al., 2000; 2006). Together, this evidence strongly suggests that an ordinal retrieval mode dominates early in sequence memory, and is subsequently bolstered by other retrieval modes such as a TCM mode. Likewise, in humans it has been shown that multi-voxel patterns from the mPFC are significantly higher for objects that share the same position information, compared to objects in different positions (Hsieh et al., 2015), suggesting convergent patterns of activation reflect shared ordinal representations within mPFC (Tiganj et al., 2017) despite the difference in object identity. Taken together, prior research and the current study show compelling evidence that the mPFC helps remember *when* events occurred by engaging an ordinal retrieval mode.

Theoretically, an ordinal retrieval mode generated by mPFC would be input to the HC during episodic memory through indirect cortical or thalamic pathways (for review see Dolleman-van der Weel et al., 2019). The engagement of an ordinal retrieval mode in HC might then allow for the rapid formation of conjunctive item-position representations (e.g., 1^st^ – A, 2^nd^ – B, etc.; Fig. 4B), and provide sequential structure without an explicit need to represent the elapsing time between items (a useful form of neural compression for temporal information). There is indirect evidence that item-position representations are reflected in rodent CA1 neurons during spatial sequence tasks (Euston and McNaughton, 2006), and direct evidence for conjunctive item-position representations during an analogous odor sequence task (Allen et al., 2016). Importantly, item-position representations are learned and retrieved *before* sequential item-item associations in non-human primates (Orlov et al., 2000). This begs the question whether episodic memories typically rely on an ordinal retrieval mode for recalling events with timelines. Subsequent replay events or other consolidation processes could either strengthen temporal context modes and/or weaken ordinal modes, although we suspect the former.

Interestingly, in monkeys ordinal transfers show graded interference over lag distances, suggesting they share similar properties with temporal context retrievals when studied in this way (Orlov et al., 2006), indicating two temporal dimensions may be complementary and normally integrated. Future neuroimaging studies in humans using ordinal transfers distributed across lag distances will be useful for examine the neural activity when these processes are both contributing to retrieval patterns.

### Temporal context retrieval modes in HC

Consistent with the literature, our results show that the HC contributes to sequence memory through the use of a temporal context retrieval mode. The HC has been shown to be important for sequence memory (Hsieh et al., 2014; Goyal et al., 2018) but its precise contribution has remained an open question. According to TCM, the HC associates items in sequences through a drifting contextual representation (Howard et al., 2005; Polyn and Kahana, 2008). As such, the presentation or retrieval of an item from a sequence elicits the retrieval of neighboring items that share a temporal context, decreasing in likelihood or strength for more distal items (Howard and Kahana, 2002). According to this framework, a temporal context retrieval mode accounts for the increased likelihood that adjacent items in word lists are recalled (Howard and Kahana, 2002). This pattern in free-recall performance predicted by temporal context memory has been validated in computational models (Howard and Kahana, 2002), behavioral studies (Kahana, 1996; Sederberg, 2010; Morton and Polyn, 2016), and neurobiological studies (Polyn and Kahana, 2008; Jenkins and Ranganath, 2010; Hsieh et al., 2014; Bladon et al., 2019). In the current sequence memory task, we reasoned that the presentation of InSeq items would elicit the retrieval of neighboring items that shared temporal contexts and lead to graded impairments in the detection of Skip probes. This is because, as the forward lag increased, the upcoming representations were less likely to be retrieved and thus less likely to interfere with an out of sequence determination. Similar patterns in performance are observed in other tasks probing temporal memory (Allen et al., 2014; 2015; DuBrow and Davachi, 2014).

The HC has shown activations consistent with the use of a TCM retrieval mode. Activation in the HC is elevated during the processing of overlapping compared to nonoverlapping sequences (Kumaran and Maguire, 2006; Brown et al., 2010; Brown and Stern, 2013). Population activity in CA1 drifts across both small- and large-time scales (Manns et al., 2007; Mankin et al., 2012; Ziv et al., 2013; Rubin et al., 2015; Mau et al., 2018). Additionally, HC lesions impair the discrimination of overlapping odor sequences (Agster et al., 2002). A recent MRI study showed that HC multi-voxel pattern similarity was higher for pairs of adjacent trials that belonged to the same temporal context within a sequence compared to pairs of sequence items that bridged between sequences, even when the temporal distance between the pairs of items was similar (Hsieh et al., 2014). The same study observed that the HC carries information about the temporal context between items within a sequence, rather than information about the objects themselves (Hsieh et al., 2014). Specifically, Hsieh et al. (2014) demonstrated that when the same sequence item is repeated, hippocampal voxel patterns were dissimilar, unless the temporal context was reinstated. Our results add to this by showing that as Skips lag further away from their InSeq location, HC activity also increases closely matching predictions of TCM.

### Limitations and theoretical considerations

While the use of specialized out of sequence probe trials provided important insight regarding different retrieval modes, and their related neurobiological substrates, several limitations of the current study remain. First, the timing of the task and self-paced design of the experiment preclude detailed item-based analyses at the neurobiological level. At the expense of repetitions of trials, future studies should include temporal jitter between sequence items to isolate signals at the different item positions. Second, similar to multivariate approaches, univariate approaches are subject to interpretational ambiguities (Hebart and Baker, 2018; Ritchie et al., 2017). However, to avoid overfitting issues, especially with time-varying representations, we chose to use a univariate approach over a multivariate approach as it is better suited to our task design to differentiate retrieval modes using different out-of-sequence probe trials.

Key questions remain concerning the interactions between ordinal and temporal context retrieval modes, and their related neurobiological constituents (mPFC and HC). While the current study did not include probe trials that investigated ordinal transfer and transpositions, future studies should examine these critical probe trial types and expect to see increased interactions between the mPFC and HC. mPFC-HC coupling may lead to conjunctive representations of temporal contexts and positional coding. As shown here, mPFC activation reflects ordinal retrieval modes (Fig. 4B) which interacts with temporal contexts in the HC (Fig. 4B). In theory, this information could be merged allowing the formation of conjunctive temporal context and item-position association in HC neurons. Direct evidence for this latter possibility was provided by recent studies that showed HC neurons encode item-context and item-position conjunctions (Komorowski et al., 2009; Allen et al., 2016), suggesting that these conjunctive representations provide the neuronal basis for time and place integrations in episodic memory. However, future studies are required for examining mPFC-HC interdependent interactions directly in humans.

## Conclusion

The results from our study highlight novel evidence that ordinal and temporal context retrieval modes both contribute to remembering items with a timeline. In particular we showed that mPFC activity reflected an ordinal retrieval mode and HC activity reflected a TCM retrieval mode. Further experiments in both animals and humans are necessary to delineate the precise mechanisms by which the mPFC engages ordinal retrievals, and how this process interacts with HC to integrate ordinality and temporal contexts in support everyday episodic memory.

## Acknowledgments

The research was conducted with funds provided by FIU to TAA and ATM, and the Feinberg Foundation to TAA. We thank Adam Kimbler for his help with coding, and Dr. Leila Allen for useful feedback on the manuscript.

